# Site-Specific and Diet-Dependent Genetic Architecture of Trabecular Bone in Male BXD Recombinant Inbred Mice

**DOI:** 10.64898/2026.11.18.767695

**Authors:** Krittikan Chanpaisaeng, Perla C. Reyes-Fernandez, Brian Dilkes, Vivian Quinlan, James C. Fleet

## Abstract

Vertebral fracture risk is influenced by trabecular bone (Tb) microarchitecture, a phenotype that is regulated by genetic and environmental factors. We examined the genetic architecture of Tb microarchitecture in the fifth lumbar vertebra (L5) and its modification by dietary calcium (Ca). Male mice from 51 BXD recombinant inbred lines were fed adequate (0.5%) or low (0.25%) Ca diets from 4–12 weeks of age. Micro-computed tomography (μCT) was used to assess seven L5 Tb phenotypes, and the response to Ca restriction (RCR) was calculated for each phenotype. Quantitative trait loci (QTL) were identified by composite interval mapping and compared with previously reported femoral Tb QTL from the same mice. Genetic background significantly influenced all L5 phenotypes and their responses to Ca restriction. We identified 127 QTL distributed across 61 loci. First, very few L5 QTL overlapped those previously identified in the femur, providing strong evidence for skeletal site-specific genetic regulation of Tb microarchitecture. Second, QTL detection was strongly dependent on dietary Ca, with only 4 of 33 loci identified under adequate Ca intake also detected under low Ca intake. Third, QTL controlling the RCR were largely distinct from those controlling L5 phenotypes under either dietary condition. Many loci affected multiple phenotypes, suggesting shared genetic regulation of Tb characteristics. Integration of genetic variation, expression QTL, and existing functional evidence identified plausible candidate genes at several high-priority loci. Together, these findings demonstrate that genetic regulation of Tb microarchitecture is both skeletal site-specific and diet-dependent, and that adaptation to dietary Ca restriction has a distinct genetic component.

## Introduction

Vertebral fractures are the most common type of osteoporotic fractures worldwide^(1^) and are associated with increased mortality^(2,3^) and reduced quality of life.^(4–7^) Because vertebrae are composed predominantly of trabecular bone, biomechanical competence depends on both trabecular bone mass and microarchitecture.^(8–11^) Consistent with this, individuals with osteoporotic fractures have markedly lower trabecular connectivity density (Conn.D) and trabecular number (Tb.N) than individuals without fractures.^(8,12,13^)

Skeletal phenotypes are under strong genetic regulation. ^(14–17^) Using data from three inbred mouse strains, Judex et al.^(18^) proposed that trabecular and cortical regions of femoral bone are under distinct genetic controls. Paternoster et al.^(19^) subsequently reported evidence for distinct genetic influences on trabecular and cortical volumetric bone mineral density (BMD) of the tibia in three human cohorts. We^(20^) and others^(21^) have also shown that trabecular and cortical bone phenotypes are differentially and independently modulated by genetics. These observations raise a related question: is genetic regulation of trabecular bone also site specific? Studies comparing trabecular bone at different skeletal sites^(22,23^), including our own comparison of the distal femur and L5 vertebra^(24^), suggest it may be.

Although these studies suggest site-specific genetic regulation of trabecular bone, most studies of its genetic architecture have focused on the femur^(20,21,25,26^) or the tibia.^(21,27^) In contrast, loci controlling vertebral trabecular bone traits have been mapped in only two studies, an F2 population of 2-month-old mice from a B6 x C3H cross^(28^), and an in silico mapping study using data from 12 inbred strains of 4-month-old mice.^(23^) In 2019 we used 63 inbred and recombinant inbred strains and found that both femur and L5 spine trabecular bone µCT phenotypes were heritable. However, the strain distribution patterns for femur and L5 spine phenotypes were divergent.^(24^) In addition, dietary calcium restriction had a greater negative impact on femur µCT traits than on L5 µCT traits. While we previously reported that low dietary calcium intake influenced the penetrance of femur µCT traits ^(20^) (i.e. gene x diet (GXD) interactions), but whether the genetic regulation of vertebral trabecular bone is similarly modified by dietary Ca remains unknown.

To directly examine the effects of genetics and GXD interactions on vertebral trabecular bone µCT phenotypes, we conducted a study using male mice from 51 BXD RI lines raised under rigorously controlled environmental conditions, with strict control of age, and exposure to either adequate or low dietary Ca. We characterized the genetic architecture of L5 trabecular bone under both dietary conditions as well as the genetic control of its response to dietary Ca restriction. Importantly, because these same mice were used in our previous analysis of femur trabecular bone^(20^), we could directly compare the loci controlling trabecular bone at two skeletal sites. We also compared our findings with other studies that identified loci controlling L5 µCT traits and vertebral BMD.^(22,23,28–30^) Our results reveal substantial differences in the loci controlling L5 and femur trabecular bone, as well as strong dependence of L5 genetic effects on dietary Ca, supporting a site-specific and diet-dependent genetic architecture of trabecular bone.

## Materials and Methods

### Experimental Design

This manuscript reports novel data that extends our early work on the impact of genetics on calcium and vitamin D metabolism as well as femur bone mass and microarchitecture.^(20,31,32^) We used a population of 51 BXD recombinant inbred (RI) mouse lines derived from C57BL/6J (B6) and DBA/2J (DBA) inbred mouse lines.^(33^) Each RI line has a unique recombination pattern of alleles from the parent lines that are fixed so that the genetic features can be replicated and studied in multiple environments for the assessment of gene x environment interactions.

Four-week-old male mice from the 51 lines were obtained from the Jackson Laboratory (Bar Harbor, ME, USA). Upon arrival mice were randomly assigned to either a 0.5% (basal) Ca or 0.25% (low) Ca diet (AIN93G base with 200 IU vitamin D_3/_kg diet, Research Diets, New Brunswick, NJ, USA) (n = 8-9 per diet group for all lines except BXD36 where n=4). The total of 809 mice were randomized to dietary groups; data for 751 mice were available for analysis (see Supplemental Table S1 for explanation of exclusions). The dietary Ca levels were chosen to meet the NRC rodent dietary Ca requirement (0.5% Ca) or to model the low level of dietary Ca intake seen in the U.S. population (0.25% Ca). ^(34^) Mice were group-housed (2-4 mice/cage) at the Purdue University animal facilities in conventional shoebox cages, maintained in rooms with UV-blocking filters over lights and a 12-hour light/dark cycle, and provided food and distilled water *ad libitum*. When the mice reached peak trabecular bone mass at 12 wks of age ^(35^) they were fasted overnight, after which the entire lumbar vertebral column was harvested, placed in 10% neutral buffered formalin for 48 h and stored in 70% ethanol afterwards. L5 vertebrae were subsequently isolated from the spinal column and muscle was gently removed. Investigators were blinded to genotype and dietary treatment, animal handling, bone sample collection, and endpoint measurements. All animal experiments complied with the ARRIVE guidelines and the Purdue Animal Care and Use Committee approved the experimental protocol.

### Micro-computed Tomography (µCT) Evaluation

L5 vertebrae were analyzed using µCT (µCT 40, Scanco Medical AG, Bassersdorf, Switzerland) with scanning parameter settings reported elsewhere.^(36^) To accommodate the large number of samples to be scanned, we executed a high-throughput scanning approach. For this, 3 vertebrae were stacked on top of each other and immobilized by inserting a blunted toothpick through the vertebral body, taking care not to damage the trabecular bone. Six stacks were placed in a 15-mm-diameter polyetherimide sample holder filled with 70% ethanol and scanned at 16-µm resolution (isotropic voxel size) using an energy level of 55 kVp, an integration time of 300 ms, and an intensity of 145 µA. Using Scanco evaluation software (version 6), each vertebra was assessed for bone volume fraction (BV/TV), trabecular number (Tb.N, mm^-1^), trabecular thickness (Tb.Th, mm), trabecular separation (Tb.Sp, mm), connectivity density (Conn.D, 1/mm^3^), structure model index (SMI), and trabecular bone tissue mineral density (Tb.TMD) as recommended elsewhere.^(35^) The region of interest was defined as 0.800 mm (50 slices) distal to the end of cranial growth plate. This region was chosen as it represents trabecular bone parameters of the entire L5 well.^(37,38^) We manually contoured trabecular bone every 10 slices with the outline 2-3 pixels away from the cortical bone, and the intermediate slices were interpolated with the contouring algorithm in the software to create a volume of interest. Morphometric parameters were evaluated using a Gaussian filter = 0.8 and a threshold of 246 (559.6 mg HA/cm^3^).

### Statistical Analysis

Statistical analysis was conducted using SAS Enterprise Guide 8.4 (SAS Institute Inc., Cary, NC). Data points with a Z-score in the extreme 2.5% of either end of a line/diet group distribution were removed as outliers. **Supplemental Table S1** reports the number of mice in each genotype and dietary treatment group for each µCT parameter after outlier removal. In addition to the data obtained from each mouse on each diet, a parameter reflecting the response to dietary Ca restriction (RCR) was calculated as the percent difference between the phenotypic value for an individual (i) fed the low Ca diet (x) and the line (j) mean for the phenotype value from the basal Ca diet (y), standardized to the line mean for the phenotypic value from the basal Ca diet and multiplied by 100, i.e., [(*x_ij_* − *ȳ_j_*) / *ȳ_j_*] ∗ 100.^(36^) Raw means for all parameters are provided in Supplemental Table S2.

For each phenotype, the covariate effect of body weight (BW) and/or femur length (FL) was determined by Pearson’s correlation and removed by linear regression when the effect on a parameter was significant.^(39^) The body size (BS)-corrected residuals were examined for normality and, when necessary, data were transformed. The complete record of co-variate corrections and transformations are provided in Supplemental Table S3. To reflect the variation among lines, Z-scores were calculated from the line means of BS-corrected residuals of the 51 BXD RI lines and their parental lines. Principal Components Analysis (PCA) was used to compare the phenotypic relationships among the seven µCT traits in the L5 spine and distal femur using femur data previously reported from the same mice.^(20^) PCA was conducted in OriginPro 2026 (OriginLab, Northampton, MA) using a Pearson’s correlation matrix. Pearson’s correlations for the 7 L5 µCT phenotypes and the three conditions (0.5% Ca, 0.25% Ca, RCR) are reported in Supplemental Table S4.

Line means of BS-corrected residuals for each dietary condition (basal, low Ca and RCR) were used for genetic mapping (**Supplemental Table S5)**. We assessed the narrow-sense heritability (*h*^2^) of each Tb phenotype using the r^2^ from a one-way Analysis of variance (ANOVA) (main effect = genotype) ^(40,41^); this was conducted separately for each diet group as well as for their RCR. Two-way ANOVA of BS-corrected residuals was used to test the main effects and interaction effects (i.e. genotype-by-diet, GXD) on each phenotype. GXD interactions were also determined by conducting one-way ANOVA to assess the impact of genetics (i.e. line) on the RCR. Using 51 BXD RI lines allowed us to reliably detect QTL accounting for 15% of the variance observed in the population with the power ∼ 0.80 and α = 0.05.^(40,41^) Using n=8/line/diet also allowed us to obtain accurate line estimates of Tb phenotypes.

### QTL Mapping

BXD genetic markers were downloaded from the GeneNetwork (https://info.genenetwork.org/species/source.php?SpeciesName=mouse&InbredSetName=BXD). For the 198 BXD strains, the file contains 7,321 markers and provides approximate locations for 10,300 recombinations (average of 52 per strain). Genetic marker locations are reported in base pairs (bp) based on the mouse genome build GRCm38/mm10. The Mouse Map Converter (MMC) tool (https://github.com/rqtl/mmconvert) was used to convert mm10 coordinates to sex-average centiMorgan (cM) values that were then used for QTL mapping analysis. After excluding uninformative genotype data from lines outside the 51 lines in our experiment, we used the findDupMarkers and drop.marker functions available in the R/qtl package (https://rqtl.org/) to remove markers with duplicate genetic locations or perfectly correlated genotypes. The final genetic map for the 51 lines contained 2,244 markers (available on request).

Composite interval mapping (CIM) was conducted on BS-corrected line means (n=51) using Windows QTL Cartographer v2.5_011 (http://statgen.ncsu.edu/qtlcart/WQTLCart.htm). Forward selection identified 5 significant background markers. CIM was carried out using a Haldane map function, 2 cM walking speed, and a 10 cM window. Phenotypes for each diet (0.5% or 0.25% Ca) group and the RCR were mapped separately. For each analysis, 1000 permutations were used to determine significance threshold in a logarithm (base 10) of odds (LOD) unit. A peak with LOD ≥ the computed permutation threshold was considered significant. Because of our interest in pleiotropy, we also tracked peaks with LOD ≥ 2, a level that many reports use to identify putative QTL.^(42^) For each significant or putative QTL, a 1.5-LOD support region, equivalent to a 95% confidence interval was defined.^(43^) To assess the extent to which genetic loci were shared between skeletal sites, L5 QTL were compared with femur trabecular bone QTL previously identified in these same mice.^(20^) L5 and femur QTL were considered overlapping when their 1.5-LOD support intervals shared a common genomic region.

### QTL Consolidation and Prioritization

All significant and putative loci for each trait and condition were combined and are reported in Supplemental Table S6. We identified many loci across all the phenotypes and multiple phenotype QTL were frequently co-localized. When this occurred, we grouped these co-localized QTL into a single locus and assigned an identification (loci ID).

Using the complete QTL loci list, we selected several loci for further investigation. We prioritized loci based on the following criteria: (1) presence of a significant QTL for a single trait (high Ca, low Ca, RCR) with an LOD > 5; (2) Significant (or one significant with one putative) QTL observed for the same phenotype in both the basal and low Ca groups and with a matched parental influence; (3) Multiple QTL observed within the same condition (≥ 3 basal, low Ca or ≥ 2 RCR) or within the low Ca group and RCR with a matched parental influence; and (4) A locus with 5 or more significant or putative QTL. We gave a score of 1 for each criterion that was met. These criteria are not mutually exclusive, e.g. a single locus may earn a score for multiple of the criteria. The loci with a score of 3 or greater were considered as *high-priority* loci that were then considered for in-depth bioinformatics analysis.

### Bioinformatic Characterization of Loci

The process we used to evaluate high-priority loci is summarized in Supplemental Fig. S1A. For each locus, single nucleotide polymorphisms (SNPs) between B6 and DBA mice were obtained using the Mouse Phenome Database (MPD, https://phenome.jax.org/snp/retrievals)^(44^) with the Sanger data set that contains data on >89 million SNP and indels from 37 inbred strains of mice.^(45^) MPD annotations (NCBI dbSNP 138) were used to categorize polymorphisms by gene attribute: i.e. intronic and non-coding, insertions/deletions, mRNA untranslated region (5’and 3’ UTR), and exon-associated (i.e. synonymous and non-synonymous codons, stop codons, splice sites, or frameshift mutations). Genes with SNPs that lead to gain or loss of a stop codon, cause a frameshift, and/or alter a splice site were automatically considered possible candidate genes. Effects of non-synonymous amino acid changes were examined for potential functional effects using PROVEAN v1.1 (score < −2.5)^(46^) and SIFT v4.0.3 (score < 0.05 ).^(47^) Genes with deleterious/damaging SNPs identified by at least one tool were considered as possible candidate genes.

Local (*cis*) expression QTL (eQTL) analysis was conducted *in silico* using publicly available microarray data from the femur mRNA of the BXD panel (GN accession: GN414) that is available at the GeneNetwork (https://info.genenetwork.org/infofile/source.php?GN_AccesionId=414). Searches were conducted on the 1.5-LOD support interval of each locus. We defined the LRS threshold as > 13.6 (∼ LOD = 3) to capture both significant and putative eQTL. Of these eQTL, we examined the correlation between the abundance of a gene transcript within the *cis* eQTL, and BS-corrected line means of each phenotype mapped to the locus. Genes with significant eQTL whose SNP is within the transcribed region of the regulated gene or within 1.5 Mb of the transcribed region (i.e. cis eQTL), and whose expression level correlated with our phenotypes (Pearson’s correlations, p < 0.05) were considered as candidate genes.

We assessed whether candidate genes are likely to have function-altering polymorphisms influencing bone microarchitecture by searching the literature and public databases for information that links the candidate genes to bone biology. The list of criteria included: (1) a significant association with bone phenotypes in GWAS; (2) a functional role in bone from gene knockout mice (from the International Mouse Phenotyping Consortium (IMPC) and the International Mouse Strain Resource (IMSR) databases; (3) evidence from animal or cell studies identifying a mechanistic role for the candidate gene in bone cell biology or other pathways known to influence bone phenotypes (from Coremine, www.coremine.com/medical/); and (4) links to gene ontology (GO) terms related to bone (from the Mouse Genome Informatics database, MGI, www.informatics.jax.org); and (5) high expression in bone cells (BioGPS, http://biogps.org/).

## Results

We previously reported the QTL influencing distal femur trabecular bone phenotypes in these same BxD mice.^(20^) We therefore used PCA to compare the relationships among the µCT traits from distal femur and L5 spine. PCA revealed substantial site-specific differences in these relationships (Fig. 1A). Although the loading vectors for femur and L5 Tb.Th were closely aligned, the vectors for most paired femur and L5 traits diverged, indicating that the relationships among trabecular phenotypes differ between the two skeletal sites. Similar differences were observed for the RCR phenotypes (Fig. 1B).

**Figure 1.**
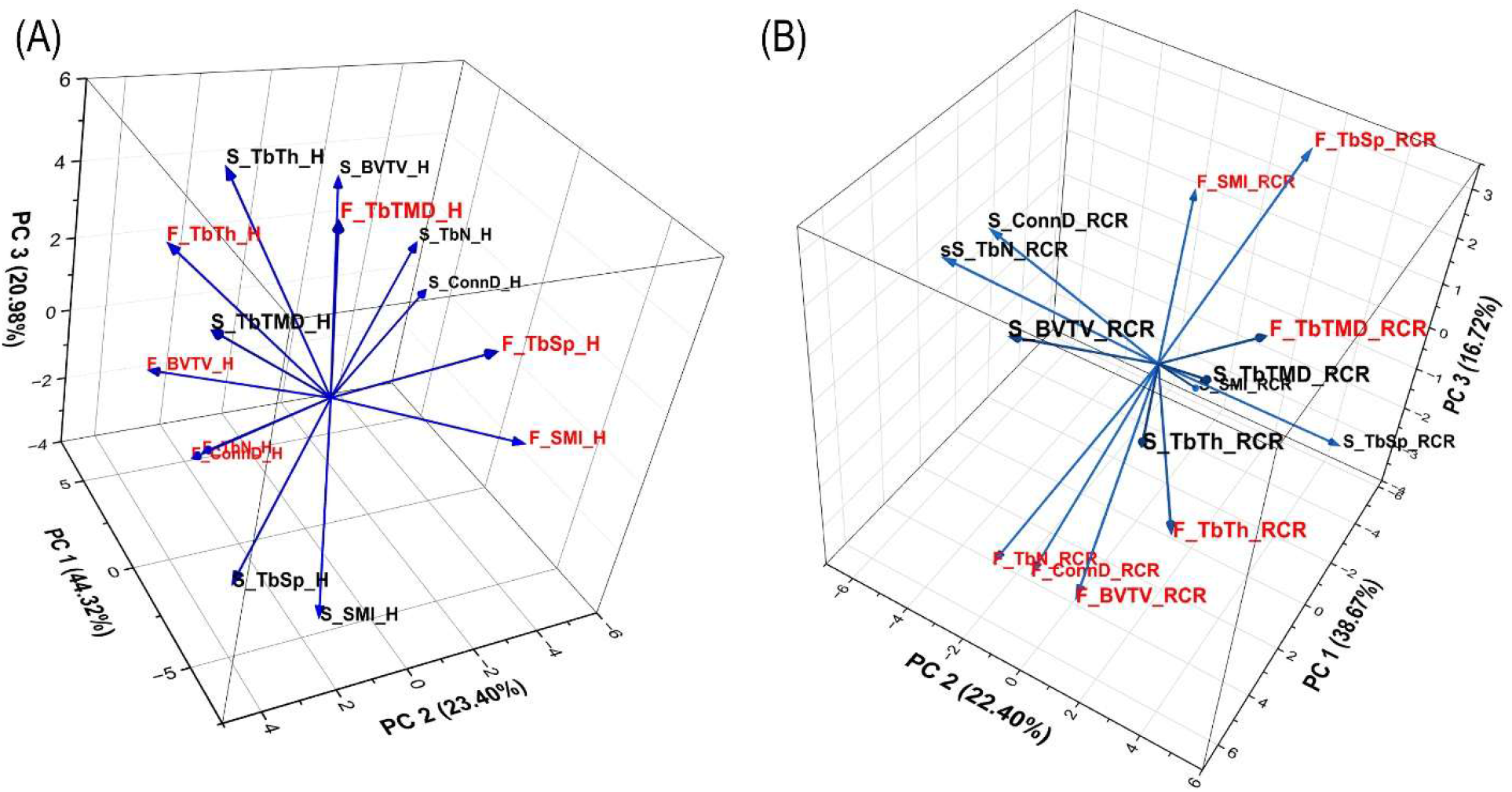
PCA Analysis Shows Divergence Between Femoral and L5 Spine trabecular bone (Tb) µCT Phenotypes in BXD RI Mouse Lines. (A) Comparison of trabecular bone µCT phenotypes in mice on a 0.5% Ca diet (H); (B) Comparison of trabecular bone µCT phenotypes for the response to calcium restriction (RCR).

In contrast, loading vectors for corresponding L5 traits measured under the 0.5% and 0.25% Ca diets were generally aligned, indicating that the overall phenotypic structure was largely preserved across dietary conditions, although some diet-dependent differences were evident (Supplemental Fig. S2A). The RCR phenotypes had a markedly different multivariate structure from the corresponding L5 phenotypes measured under either dietary condition (Supplemental Fig. S2 B,C), indicating that variation in the physiological response to Ca restriction captures a distinct dimension of trabecular bone variation. Strain distribution patterns for the μCT phenotypes were consistent with the PCA results (Fig. 2, Supplemental Fig. S3). For example, the strain distribution patterns for L5 Tb.Th were similar between the 0.5% and 0.25% Ca diets, whereas they differed markedly between L5 Tb.Th and its RCR (Fig. 2B) and between femur and L5 Tb.Th on the 0.5% Ca diet (Fig. 2C).

**Figure 2.**
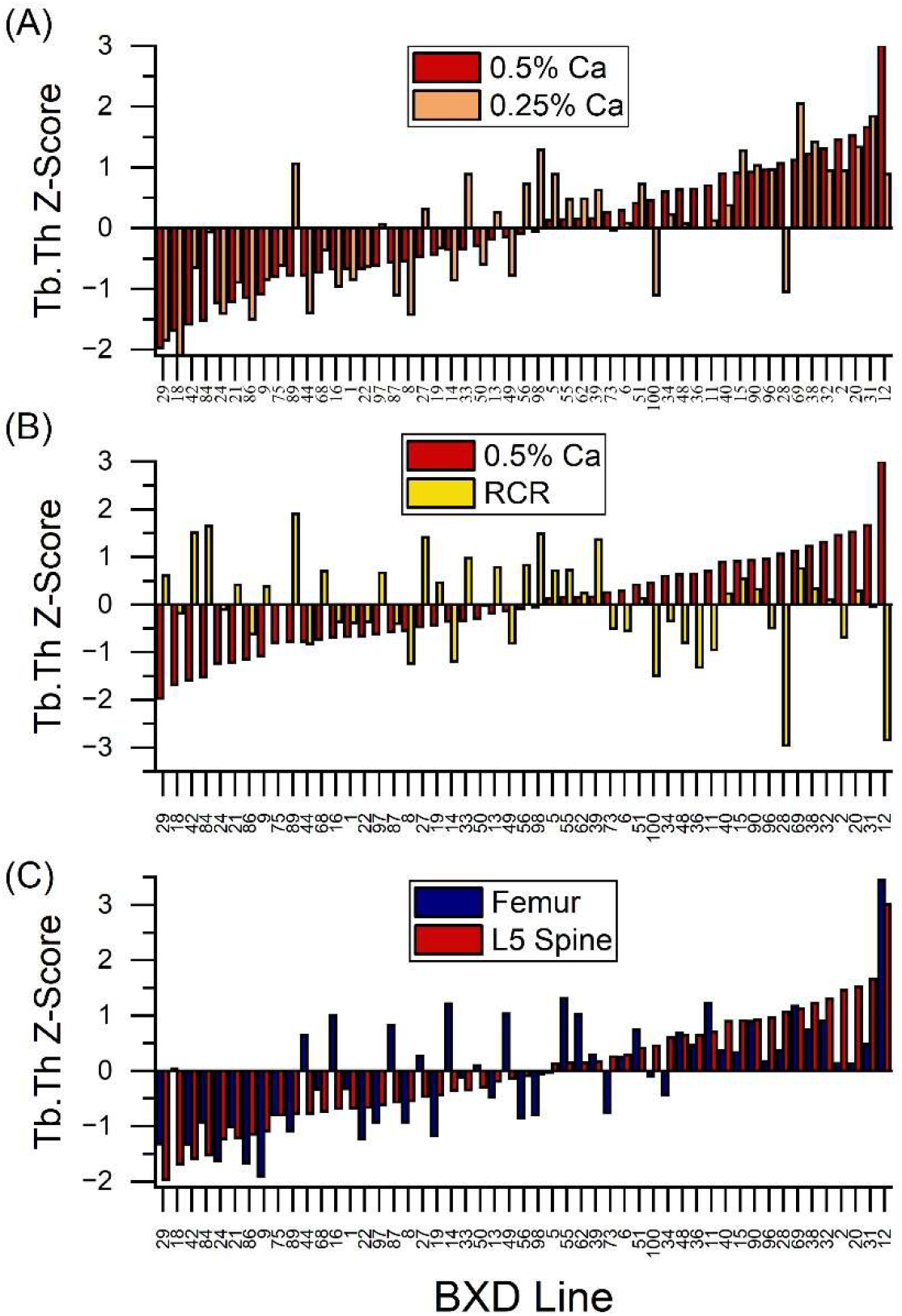
Strain distribution patterns for trabecular thickness (Tb.Th) across the 51 BXD lines. (A) Comparison of the distribution of L5 Tb.Th for lines on the 0.5% Ca diet (red bars, plotted low to high) to mice on the 0.25% Ca diet (orange bars); (B) Comparison of the distribution of L5 Tb.Th for lines on the 0.5% Ca diet (red bars, plotted low to high) to the line response to dietary calcium restriction (RCR, yellow bars); (C) Comparison of the distribution of L5 Tb.Th (red bar, plotted low to high) to femur Tb.Th (blue bars) for lines on the 0.5% Ca diet. For each plot, the values are reported as phenotype Z scores.

ANOVA revealed significant line effects on each L5 µCT phenotype, i.e. genetic (Table 1). Heritability estimates for L5 µCT traits ranged from 0.36 to 0.63 in mice fed the 0.5% Ca diet and from 0.32 to 0.64 in mice fed the 0.25% Ca diet (Table 2). Diet significantly affected all L5 µCT phenotypes, although only Conn.D showed a significant Line x Diet interaction. However, line significantly affected the RCR for each phenotype, demonstrating genetic variation in the response to dietary Ca restriction. Heritability estimates for the RCR phenotypes were more modest (0.23-0.35), similar to what we previously reported for femur trabecular bone RCR µCT phenotypes.^(20^)

**Table 1.**
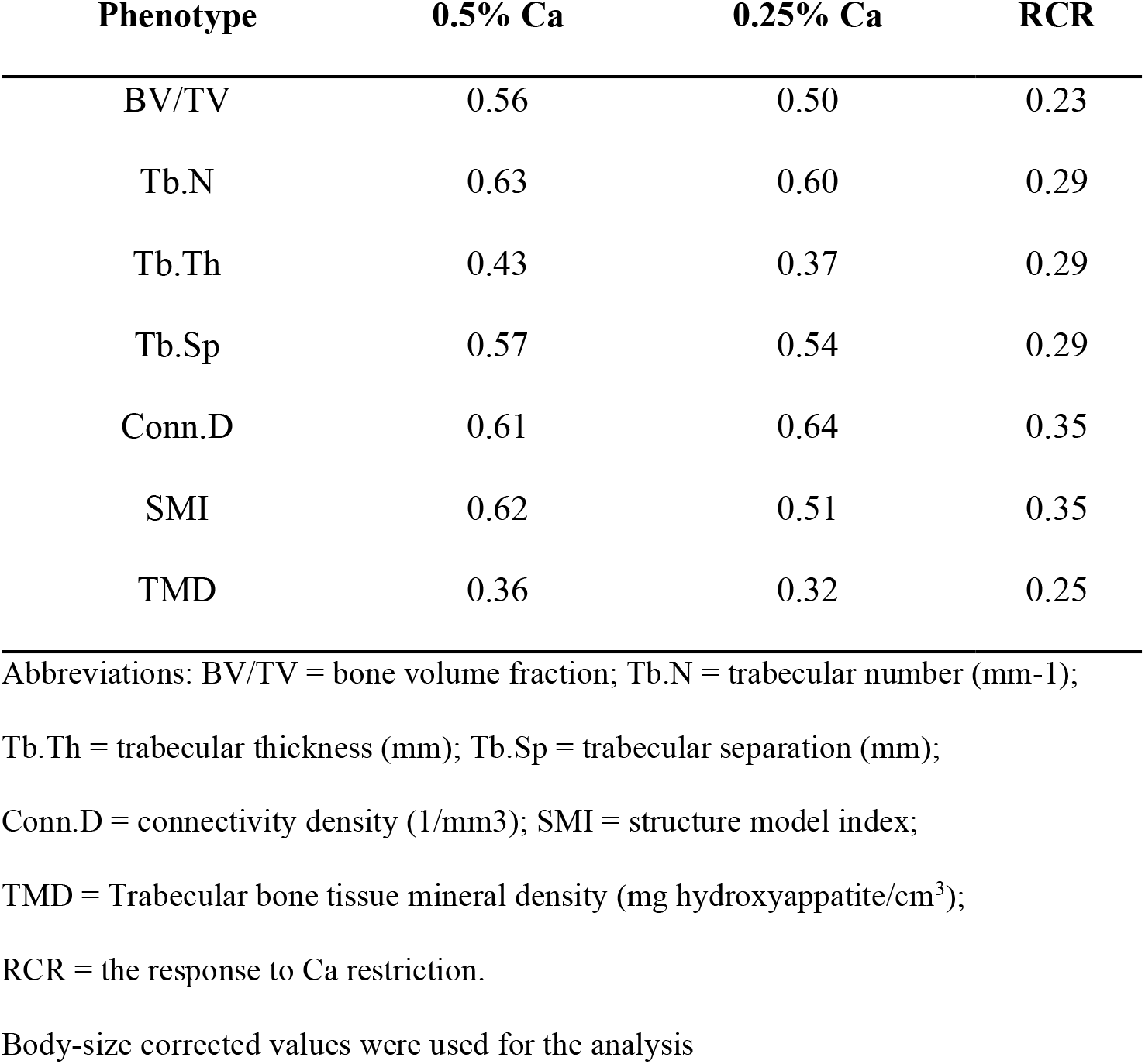
Heritability Estimates (h^2^) for L5 Trabecular Bone (Tb) Traits for Each Dietary Environment and for Their Response to Calcium Restriction (RCR)

**Table 2.** Summary of candidate genes for each of the 10 high-priority loci.

| QTL ID | Type of Variant<sup>1</sup> | Gene 2 | Type<sup>2</sup> | Candidate Rank<sup>3</sup> | L5 Phenotype<sup>4</sup> |
|---|---|---|---|---|---|
| 1A | eQTL | Kcnq5 | 5, 7, 8 | 2 | ConnD (H, B, 3.75/6.68), BVTV (H, B, 5.25), SMI (H, D, 7.76) |
| 1E | Cn | Ephx1 | 2 | 2 | ConnD (H, B, 5.20), ConnD (L, B, 3.13), TbN (RCR, D, 4.38), TbSp (RCR, B, 5.35), TbSp (H, D, 4.10), SMI (L, D, 2.93) |
|  | Cn | Lyplal1 | 1,5 | 4 |  |
|  | Cn | Mia3 | 2 | 2 |  |
| 2E | Cn | Ccn5/Wisp2 | 2 | 4 | TbSp (L, B, 3.62), TbSp (H, B, 5.20), ConnD (L, D, 4.56), TbN (L, D, 5.40), BVTV (L, D, 3.04), SMI (L, B, 3.96) |
|  | Cn | Cd40 | 1,4 | 1 |  |
|  | Cn | Plcg1 | 2,5 | 1 |  |
|  | eQTL | Rnf114/Zpf313 | 2,7 | 2 |  |
| 3D | Cf | Dkk2 | 1,4 | 1 | TbN (H, D, 2.33), BVTV (L, D, 3.98), ConnD (L, D, 2.12), TbSp (H, B, 2.93), SMI (RCR, D, 5.35) |
|  | Cn | Pdlim5 | 1,5,6 | 1 |  |
|  | Cn | Sec24d | 1,4,5 | 1 |  |
| 5A | eQTL | Cacna2d1 | 2,7, 8 | 2 | BVTV (H, D, 6.84), SMI (H, B, 6.18) |
| 7A | Cn, StopG | Cd33 (Siglec-3) | 1,3,4 | 1 | BVTV (L, D, 2.27/3.26), SMI (L, B, 3.6/7.45) |
|  | eQTL | Lin37 | 7 | 4 |  |
| 7B | Cn | Lrrk1 | 1,6 | 3 | TMD (RCR, B, 3.12), SMI (RCR, D, 5.4/5.77), TbSp (RCR, D, 3.81), BVTV (RCR, B, 2.65) |
|  | Cn | Synm | 2,4 | 1 |  |
|  | Cn | Pcsk6 | NA | 5 |  |
| 12C | Cn | Kif26a | 6 | 4 | ConnD (L, D, 3.71/8.71), ConnD (H, D, 7.04), TbTh (L, B, 2.47), TbN (L, D, 2.32), TbN (H, D, 7.57), BVTV (L, D, 2.15), TbSp (L, B, 6.92), TbSp (H, B, 6.56), TMD (RCR, D, 2.14) |
|  | Cn | Syne3 | 5 | 3 |  |
|  | eQTL | Ppp1r13b | 2,5,7 | 1 |  |
|  | eQTL, Cn | Klc1 | 2,5,7 | 1 |  |
|  | eQTL, SSA | Cdc42bpb | 5,7 | 4 |  |
| 15A | Cn | Dnah5 | 8 | 5 | SMI (L, D, 9.40), ConnD (H, B, 2.42), BVTV (L, B, 6.10), SMI (RCR, B, 2.20) |
| 20A | Cn | Vgll1 | 8 | 5 | TbN (RCR, B, 5.72), ConnD (RCR, B, 8.58) |
|  | Cn | Zfp36l3 | 8 | 5 |  |

<sup>1</sup> Cn,  non-synonymous; Cf, frameshift; Stop G, Stop codon gained; SSA, Splice site added;  eQTL, expression QTL in bone
<sup>2</sup> Type: 1 = match to >= 3 annotations; 2 = match to >= 2 annotations; 3 = Stop Gain or Loss; 4= bone phenotype in KO mouse;
5 = in a human bone GWAS loci; 6 = in osteocyte transcript signature or bone ignorome;
7 = bone eQTL that correlates with L5 phenotypes; 8 = only viable candidate at locus
<sup>3</sup> Rank: 1 = highest, 5 = lowest
<sup>4</sup> L5 spine uCT phenotypes; H = 0.5% Ca diet; L  = 0.25% Ca diet; RCR = response to calcium restriction;
B = C57/BL6; D = DBA/2J; Values = LOD scores at the locus for a phenotype

We identified 127 individual QTL (53 significant, 74 putative) distributed across 61 loci (Fig. 3). The complete list of L5 QTL, and their relationship to our previous QTL mapping for other bone- and mineral-related traits^(20,31,32^), is provided in Supplemental Fig. S6. Very few L5 loci overlapped with the femoral trabecular bone QTL previously identified in these same mice^(20^), demonstrating substantial differences in the genetic architecture of trabecular bone between the two skeletal sites. Genetic effects on L5 trabecular bone were also strongly dependent on dietary Ca. Whereas 14 pairs of femur trabecular bone QTL were detected under both Ca conditions at 10 loci^(20^), only 6 such pairs were observed for L5 traits, at 4 loci. Most L5 QTL were detected only under one dietary condition, providing evidence for diet-dependent expression, or penetrance, of genetic effects on vertebral trabecular bone. This diet dependence is illustrated by Conn.D (Fig. 4, Supplemental Fig. S4). QTL on chromosomes 6 and 12 were detected under both dietary conditions, whereas QTL on chromosomes 1 and 19 were detected only under 0.5% Ca and a chromosome 2 QTL was detected only with 0.25% Ca (Fig 4A,B). The genetic architecture of the response to Ca restriction was also largely distinct from that of trabecular bone under low-Ca conditions. Only 2 of 34 RCR QTL co-localized with a QTL detected in mice fed 0.25% Ca (e.g. Conn.D, Fig. 4C).

**Figure 3.**
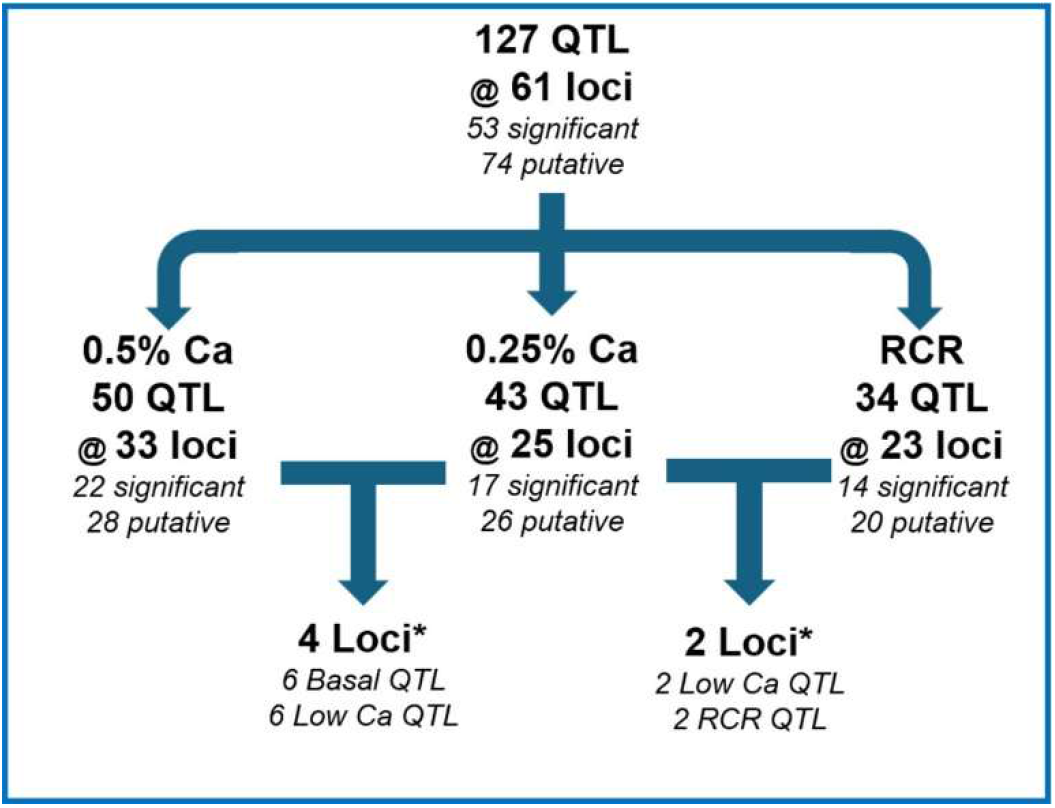
A summary of total and significant QTL affecting L5 spine μCT parameters. Loci are provided for each dietary calcium (Ca) group and for the RCR parameter. Values are presented as total QTL, significant loci (i.e., those that met the permutation threshold), and putative loci (LOD ≥ 2.0). *Overlap between loci in groups. Note: Only QTL where the same trait was seen at the same loci under both diet conditions are reported. The significant overlap count requires that at least one of the two QTL for a phenotype be significant.

**Figure 4.**
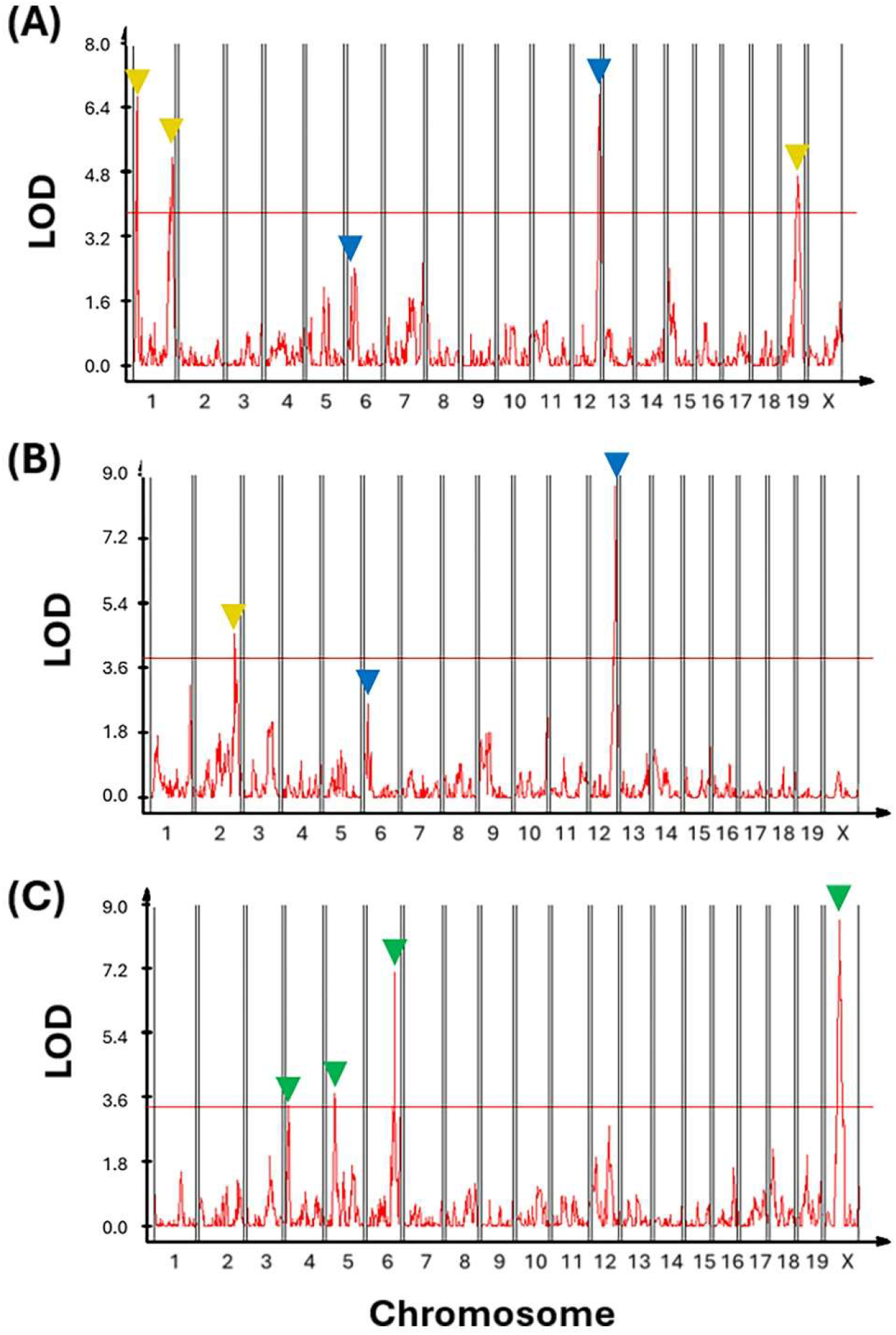
QTL Plots for Connectivity Density (Conn.D) for BXD lines fed (A) 0.5% Ca diet, (B) 0.25% Ca diet, or (C) the response to dietary Ca restriction (RCR). Lines in each plot represent the permutation threshold for significance. Yellow triangles are loci that are unique to a dietary condition, blue triangles are loci that are significant or putative (LOD>2) in both dietary conditions, green triangles are loci that are unique to the RCR.

Thirty of the 61 total loci we identified contained QTL for multiple µCT phenotypes (Fig. 5). These multi-trait loci are consistent with shared genetic effects on trabecular phenotypes and may reflect genetic control of an underlying biological property, or latent trait, that controls multiple phenotypes. Ten multi-trait loci were identified as high-priority sites. For example, the chromosome 12 locus between 97.7 and 112.9 Mb contained 10 QTL affecting six µCT traits, including QTL for Conn.D, Tb.Sp, and Tb.N that were detected under both dietary conditions. In addition, three high-priority loci contained QTL for two or more RCR phenotypes, e.g. the chromosome 7 locus contained QTL for the RCR of TMD, SMI, Tb.Sp, and BV/TV.

**Figure 5.**
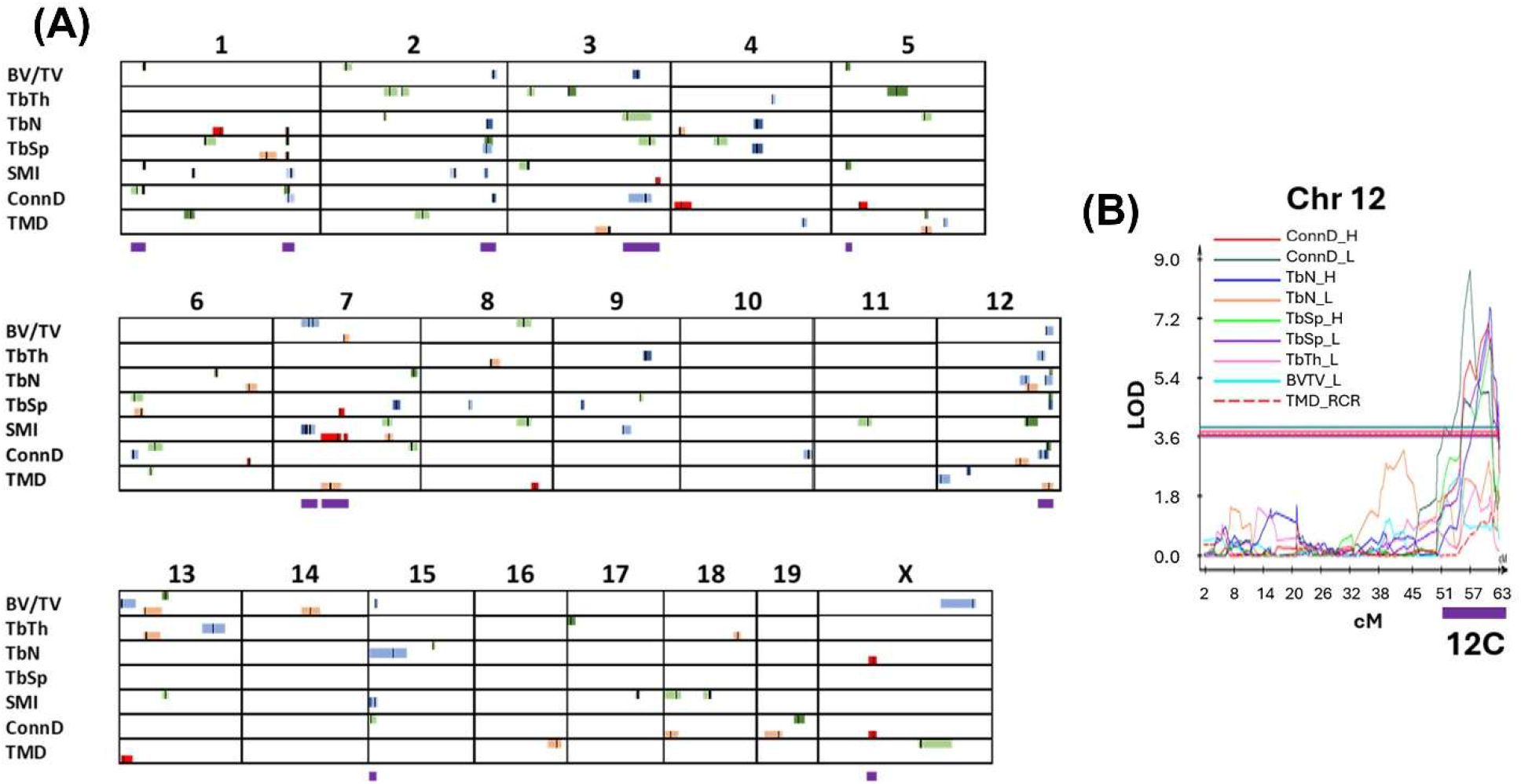
(A) A summary of QTL identified for all 7 L5 spine trabecular bone traits in the 0.5% Ca or 0.25% Ca diet groups or in the response to calcium restriction (RCR). The confidence interval for each QTL is represented as a colored bar (green = 0.5% Ca, blue = 0.25% Ca, and red = RCR) and the peak location of each QTL is shown as a vertical black line within the bar. Putative QTLs are shown as a lighter shade of each group color. The 10 high-priority, multi-trait loci chosen for bioinformatic analysis are identified by purple bars at the bottom of a chromosome. (B) The line plot for the 9 individual phenotype QTLs in the 12C locus. The horizontal lines are the permutation threshold values for each of the traits. The purple bar under the X axis is the 1.5-LOD region for loci 12C.

To investigate potential biological mechanisms underlying the multi-trait loci, we examined the 1.5-LOD support regions for the 10 high-priority loci and identified many polymorphisms affecting a large number of genes (supplemental Table S7, S10). We prioritized genes with evidence linking them to bone or bone cell biology and identified plausible candidate genes within eight of the ten high-priority loci (Table 3, Supplemental Table S11). These included genes previously shown to cause bone phenotypes when deleted in mice (Cd40, Dkk2, Sec24d, Cd33, Synm), genes linked to human GWAS controlling a bone phenotype (Kcnq5, Lyplal1, Plcg1, Sec24d, Pdim5, Tjp1, Ppplr13b, Klc1, Cdc42bpb), and those in the osteocyte transcript signature^(48^) (Lrrk1, Kif26a) or in the bone “Ignorome”^(21^)(Pdlim5).

## Discussion

A major finding of our study is that the genetic architecture of trabecular bone differs between two anatomical sites. Very few loci controlling L5 trabecular bone overlapped those controlling femoral trabecular bone^(20^) despite the two skeletal sites being examined under the same genetic and environmental conditions. Consistent with this genetic divergence, PCA also revealed substantial site-specific differences in the multivariate structure of the trabecular bone phenotypes (Fig. 1A). Kemp et al.^(14^) previously reported that BMD-associated variants may exert site-specific effects that could expand our understanding of how bone biology is regulated. Our data extend this observation by demonstrating site-specific genetic control of trabecular microarchitecture. The biological basis for these site-specific genetic effects on trabecular bone is not known. Possible explanations include differences in the mechanical loading environments at the femur and vertebrae or site-specific differences in bone cell function.^(48–50^) Such differences could cause the same genetic variant to have different effects on trabecular bone at different skeletal sites. Our findings indicate that researchers should not assume that genetic variation controlling bone biology in one site applies equally to other sites.

Our study advances our understanding of the genetics controlling L5 spine microarchitecture in other important ways. First, with its extended panel of µCT phenotypes and improved marker density, our study reveals many new loci that were not previously reported by others. Second, our data show that the dietary environment has a significant impact on the detection of L5 µCT loci with only 4 of the 33 loci controlling basal L5 µCT phenotypes also detected in mice fed the low Ca diet. These findings are consistent with our previous reports showing that the genetic regulation of femoral bone phenotypes and mineral metabolism are affected by low dietary Ca intake.^(20,31,32^) Evidence for diet-dependent QTL detection was also reported by Carson et al. ^(51^), who found that 20% of QTL controlling trabecular bone phenotypes were detected only when mice were fed low-fat diets. Third, by using recombinant inbred lines, we identified loci controlling the physiological adaptation of L5 µCT phenotypes to inadequate dietary Ca (i.e. the response to Ca restriction parameter), most of which were distinct from loci controlling L5 phenotypes under either dietary condition. Together, these findings demonstrate that dietary Ca does not simply alter L5 trabecular bone phenotypes but modifies the genetic effects underlying them and reveals distinct genetic control of the physiological response to Ca restriction.

An advantage of forward genetics is that multi-trait loci identified through natural genetic variation can reveal genes and biological mechanisms that coordinately influence multiple aspects of bone biology^(52^)(see supplemental Table S11). The most interesting locus we identified was on chromosome 12 (97.8-112.9 Mb), a locus Bouxsein et al.^(28^) previously identified using B6C3-F2 mice. We found this locus contained 10 QTL affecting 6 different traits: Tb.Th (low Ca diet), Tb.N (both diets), Conn.D (both diets), Tb.Sp (both diets), BV/TV (low Ca diet), and TMD (RCR). Within this locus, we found three potential candidate genes. The Klc1 (kinesin light chain 1) gene contains polymorphisms affecting both coding and gene expression. Klc1 is an adapter protein mediating vesicular transport along microtubules, a process required by osteoclasts during differentiation and resorption.^(53^) Klc1 mRNA is negatively correlated with BV/TV, Tb.N, and Conn.D (Supplemental Table S9) and GWAS analysis in the MSK-KP^(54^) shows that variation near Klc1 controls estimated BMD from heel ultrasound in humans (eBMD; p=1.31 x E-31).

Ppp1r13b (Trp53bp2 protein phosphatase 1 regulator subunit 13b) increases transcription of pro-apoptotic target genes by p53^(55^), a known negative regulator of osteoblast differentiation, bone formation, and osteoblast-driven osteoclastogenesis.^(56^) It has cis-eQTL-associated variation in expression where the B6 allele increases Ppp1r13b mRNA and is associated with lower BV/TV, Tb.N, and Conn.D (Supplemental Table S9). In addition, variation near Ppp1r13b in humans is a GWAS hit for eBMD (p=1.6E-13). Finally, the Kif26a (Kinesin family member 26A) gene contains a non-synonymous coding variant, and it encodes a non-traditional kinesin that affects cell positioning and migration by regulating cytoskeletal organization and cell signaling. Its paralog Kif26b can promote osteoblast differentiation.^(57^) Kif26 family members are degraded in response to Wnt5A-ROR signaling, a pathway that promotes osteoclastogenesis and bone resorption,^(58^) suggesting Kif26a could antagonize these processes. Although none of these three genes have been extensively studied in bone biology, their genetic and biological associations make them plausible candidates for mediating the multi-trait effects of the chromosome 12 locus. The high-priority chromosome 7 locus (49.8-75.9 Mb) that controls RCR phenotypes for TMD, BV/TV, SMI, and Tb.Sp contains a non-synonymous variant in *Lrrk1,* a gene highly expressed in late-stage osteoclasts.^(59^) Lrrk1 regulates osteoclast function through the small GTPases RAC1 and CDC42^(60,61^) and loss of Lrrk1 causes osteopetrosis in humans and mice because of impaired osteoclast-mediated bone resorption.^(59,62,63^) In mice exposed to low dietary Ca, the B6 allele at this locus reduced the loss of BV/TV and TMD, prevented increased Tb.Sp, and maintained a SMI that reflects stronger bone. This coordinated response is consistent with reduced LRRK1-dependent osteoclast activity attenuating bone loss during dietary Ca stress, although the functional effect of the B6 Lrrk1 variant must be experimentally established. Interestingly, Lrrk1 is also included in the osteocyte transcriptome signature identified by Youlten et al. ^(48^) suggesting that its skeletal effects may not be restricted to osteoclasts.

Our study has several strengths. First, while several studies have shown genetics regulates vertebral bone^(22,23,28–30,64^), only two examined L5 spine µCT traits. One mapped QTL in 912 four-month-old female F2 C57BL/6j x C3H/HeJ mice using a sparse marker map,^(28^) while the other conducted an *in silico* genome scan in 4-month-old male mice from 12 inbred strains.^(23^) Our analysis of the L5 spine in male mice from 51 BXD RI lines both confirms and extends this earlier work. We found loci on some of the same chromosomes as these earlier reports (e.g. BV/TV, Tb.N, Tb.Th on Chr 12, Supplemental Table S6) as well as loci not previously reported. Differences between the three L5 studies likely reflect sex-dependent genetic effects,^(65^) polymorphisms specific to each population^(45^), and differences in mapping design and resolution. The denser BXD genotype map and accumulated recombination in the RI panel also provide greater mapping resolution than the earlier studies. Second, we strictly controlled all the environmental factors that could influence bone, including the use of defined diets. Our two-diet design allowed us to distinguish loci controlling trabecular bone phenotypes that were highly penetrant across diets from those that are sensitive to low dietary Ca intake, including diet-sensitive effects specific to the L5 vertebrae.

Nonetheless, our study has certain limitations. First, our study was designed to capture genetic modulation of trabecular bone phenotypes and our candidate gene analysis prioritized genes with known roles in bone cell biology. However, bone is also regulated by crosstalk with multiple tissues, including kidney, intestine, and muscle. Thus, our candidate gene assessment is not comprehensive. Second, we used a femur mRNA dataset from BXD mice for our eQTL analysis, assuming that genetically regulated bone-cell activity is similar between the L5 vertebra and femur. However, our identification of site-specific genetic regulation of trabecular bone phenotypes indicates that femoral eQTLs may not accurately represent regulatory effects in L5 vertebrae. Third, while our study has improved mapping resolution compared to the earlier published papers, its resolution remains limited by the number of RI lines and the underlying haplotype structure and LD of the panel. Consequently, many loci encompass multiple genes and regulatory variants, limiting our ability to definitively identify causal genes and variants. Lastly, our study only used male mice and therefore may not capture genetic effects that are sex dependent or influenced by female sex hormones.

## Conclusion

This study is the first to use the BXD RI panel to map genetic loci controlling L5 trabecular bone phenotypes and their response to dietary Ca restriction. Our major finding is that the genetic architecture of trabecular bone traits differs substantially between skeletal sites, with only modest overlap between L5 spine and the distal femur QTL in the same mice. In addition, we extended our earlier observations of GXD interactions affecting vertebral trabecular bone phenotypes^(24^) by identifying loci underlying these interactions, including loci controlling the physiological response to dietary Ca restriction that were largely distinct from those controlling basal phenotypes. Bioinformatics analysis identified promising candidate genes for several loci (Supplemental Table S9), providing targets for future mechanistic studies. Together, our findings demonstrate that genetic regulation of trabecular bone microarchitecture depends on both skeletal site and dietary environment.

## Acknowledgments

This work was supported by NIH awards DK118036 and ES019103 to JCF, by start-up funds to JCF at the University of Texas at Austin, and with support from the Purdue Bone and Body Composition Core within the Indiana Clinical and Translational Sciences Institute (Award Number UL1TR001108 from the NIH, National Center for Advancing Translational Sciences, Clinical and Translational Sciences Award). KC was supported by a Royal Thai Government Scholarship while conducting some of this work as a PhD candidate at Purdue University. PRF was supported by a scholarship from CONACyT Mexico while conducting some of this work as a PhD candidate at Purdue University. We acknowledge Dr. Rebecca M. Replogle for conducting the animal experiment, and initial sample collection, and Ms. Sarah Main who assisted with the µCT analysis of the L5 spine samples.

## Author’s Roles

<u>Conceptualization</u>: JCF; <u>Data curation</u>: KC, PRF, and JCF; <u>Formal analysis</u>: JCF, BD, VQ, and KC; <u>Funding acquisition</u>: JCF; <u>Investigation</u>: JCF, PRF, KC, and VQ; <u>Methodology</u>: JCF, KC, PRF, and VQ; <u>Project Administration</u>: JCF; <u>Resources</u>: JCF, PRF, KC; <u>Validation</u>: KC, BD, and JCF; <u>Visualization</u>: KC, BD, and JCF; <u>Writing – Original Draft Preparation</u>: KC, BD, and JCF; <u>Writing – Review & Editing</u>: KC, PRF, BD, VQ, and JCF. JCF takes responsibility for the integrity of the data analysis.

## Data Availability

Unadjusted and covariate adjusted values for all phenotypes of all BXD RI lines have been provided as supplemental table S2 and S3. Individual animal level data are available upon request. BXD genotyping data is available at www.genenetwork.org/webqtl/main.py?FormID=sharinginfo&GN_AccessionId=600. QTL results are available as Supplemental Table S5 and Supplemental Fig. S4.

## Funding Statement

This work was supported by NIH award ES019103 to JCF and from the Purdue Bone and Body Composition Core within the Indiana Clinical and Translational Sciences Institute fund (Award Number UL1TR001108 from the NIH, National Center for Advancing Translational Sciences, Clinical and Translational Sciences Award). KC was supported by a Royal Thai Government Scholarship while conducting some of this work as a PhD candidate at Purdue University. PRF was supported by a scholarship from CONACyT Mexico while conducting some of this work as a PhD candidate at Purdue University.

The Authors have no conflicts to report.

## Data availability

Line averaged raw data and co-variate corrected, transformed data is available in the preliminary data. Data for individual mice is available upon request.

**Supplemental data** are included with the submission.

## Notes

### Competing Interest Statement

The authors have declared no competing interest.

