## Supplemental figures for "Site-Specific and Diet-Dependent Genetic Architecture of Trabecular Bone in Male BXD Recombinant Inbred Mice"

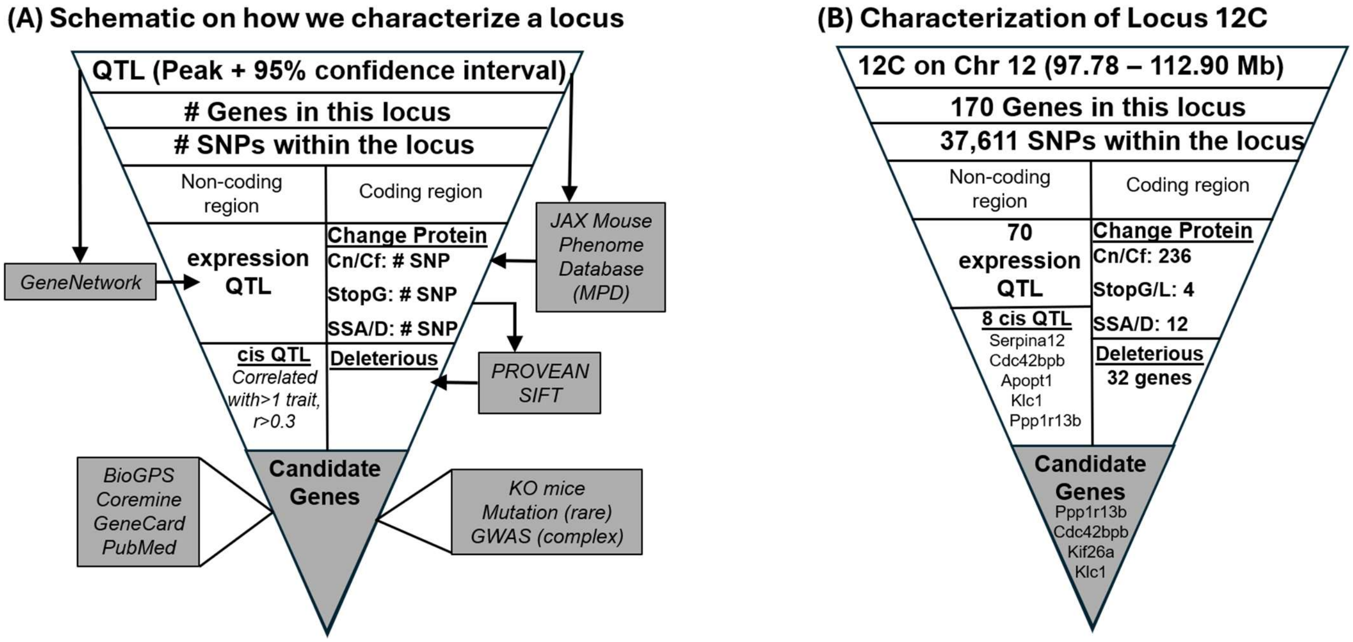


**Supplemental Figure S1. Procedure for filtering polymorphisms within a loci to identify candidate genes.** (A) General schematic identifying tools used for filtering polymorphisms within a QTL. (B) Specific application of the procedure to QTL 12 C on Chromosome 12.

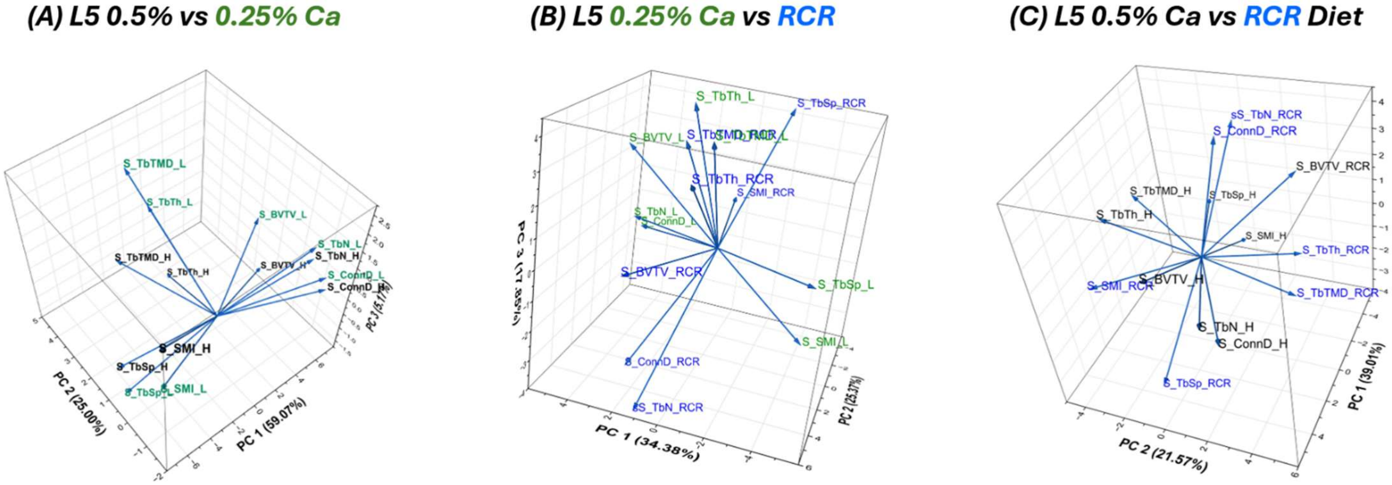


**Supplemental Figure S2. Principal Components Analysis of L5 spine microCT phenotypes.** (A) Data from BXD mice fed either a diet containing 0.5% or 0.25% diets. (B) Data from BXD mice fed the 0.25% calcium diet vs the response of mice to the low calcium diet (RCR). (C) Data from BXD mice fed the 0.5% calcium diet vs the response of mice to the low calcium diet. Mean data by BXD line was used (n=4-8 mice/line/diet).

**Figure S3. Strain distribution patterns comparing L5 Spine microCT phenotypes under three conditions (low 0.25% vs high 0.5% calcium diet; low calcium diet vs the response to low dietary calcium (RCR); high calcium diet vs RCR.** (A) BV/TV, (B) Tb.Th, (C) Tb.N, (D) Tb.Sp, (E) SMI, (F) Conn.D, (G) TMD

### (A) L5 Spine BV/TV

#### Low vs High Ca Diet

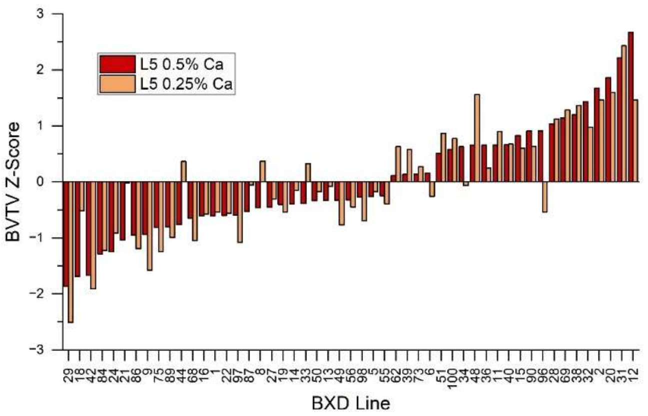


#### Low Ca Diet vs RCR

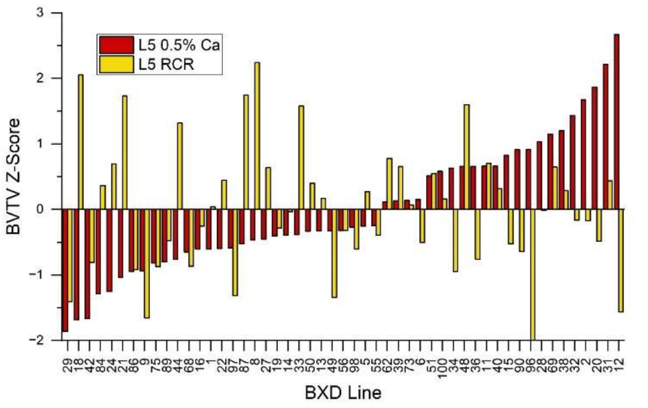


#### L5 spine vs Femur

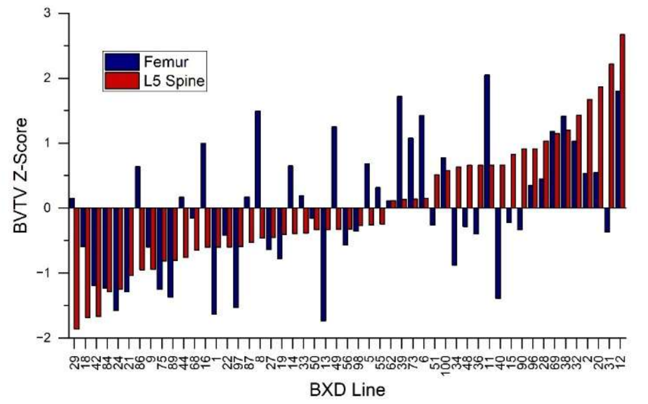


## (B) L5 Tb.Th

#### Low vs High Ca Diet

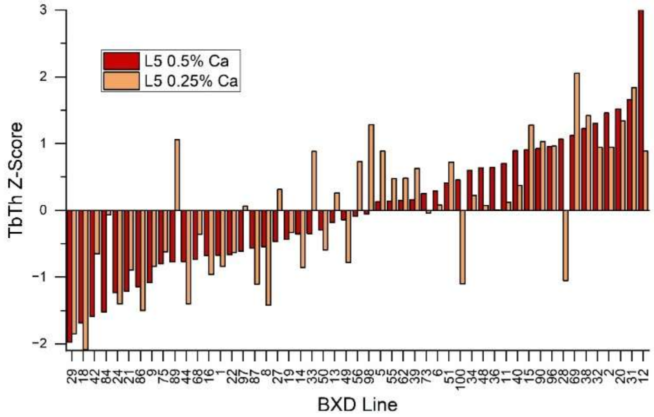


#### Low Ca Diet vs RCR

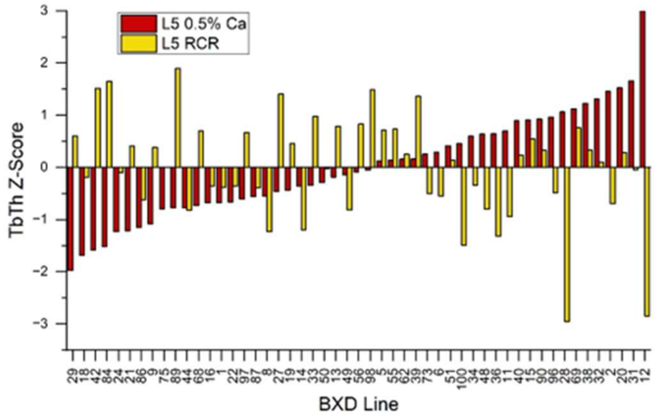


#### L5 spine vs Femur

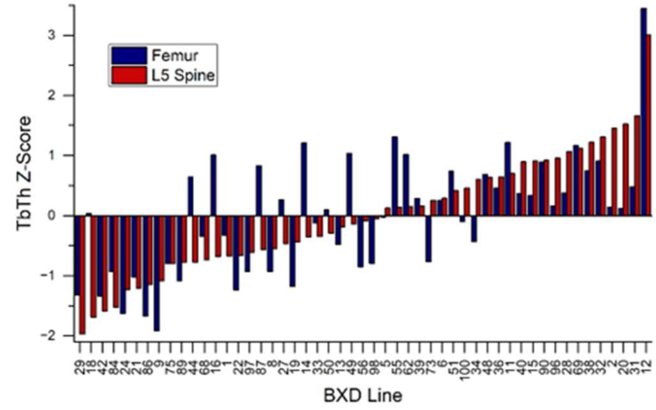

## (C) L5 Tb.N

#### Low vs High Ca Diet

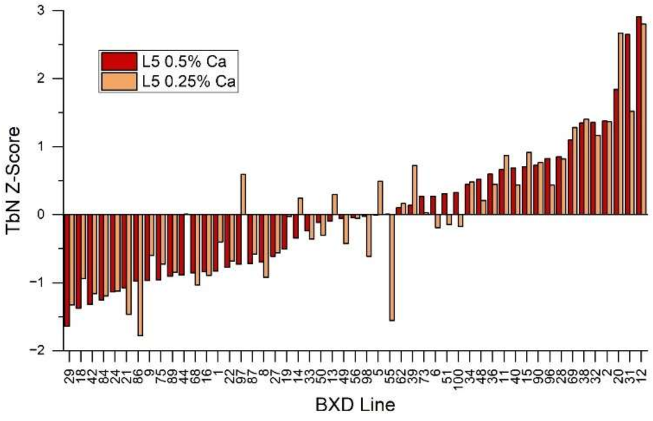


#### Low Ca Diet vs RCR

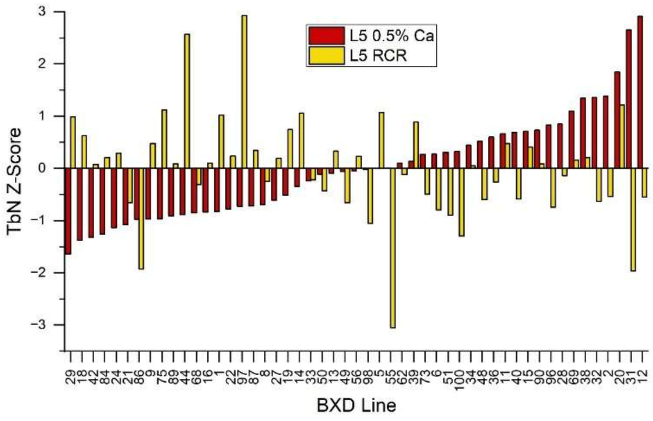


#### L5 spine vs Femur

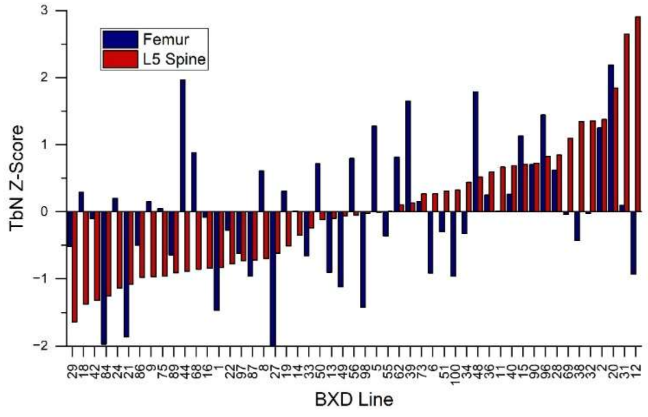


## (D) L5 Tb.Sp

#### Low vs High Ca Diet

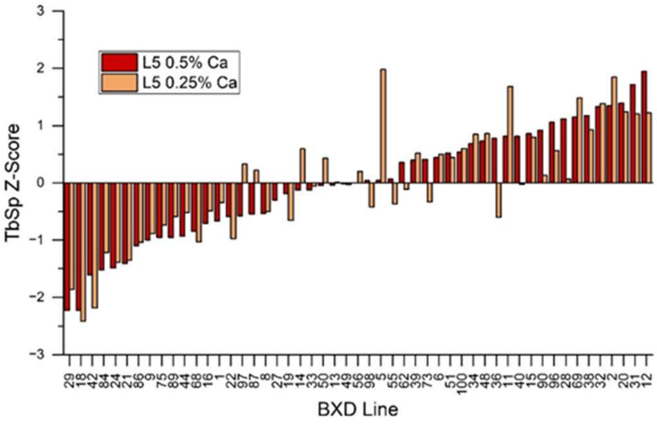


#### Low Ca Diet vs RCR

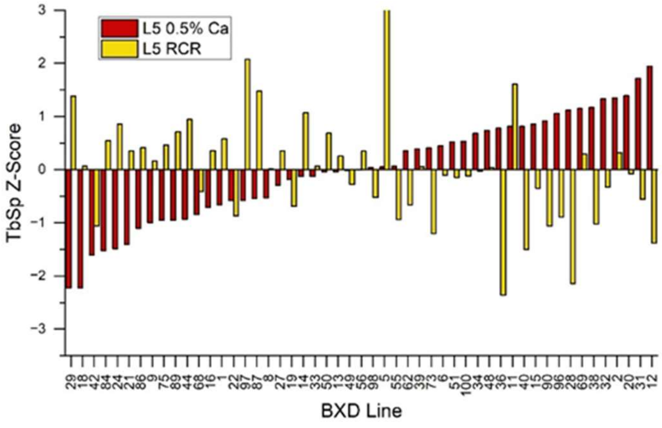


#### L5 spine vs Femur

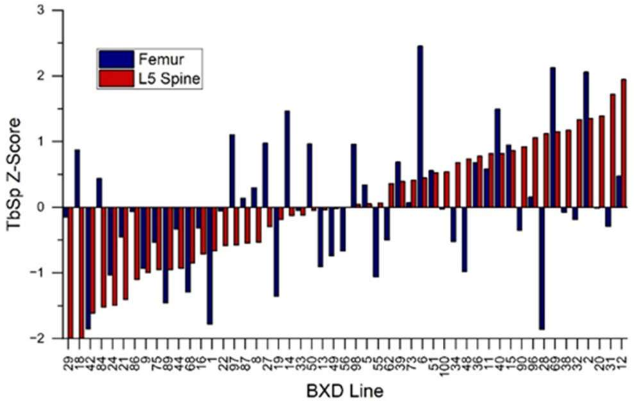

**(E) L5 SMI**
**Low vs High Ca Diet**

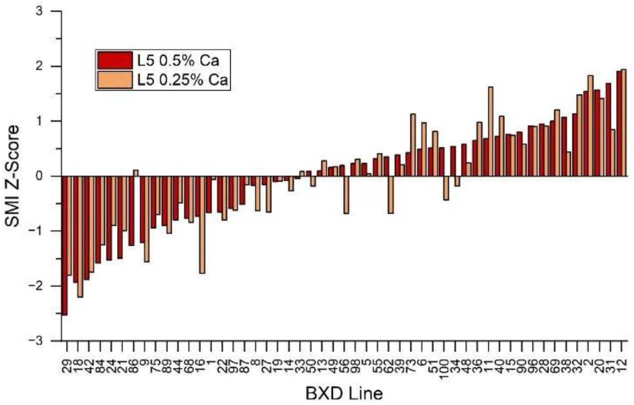


**(F) L5 Conn.D**
**Low vs High Ca Diet**

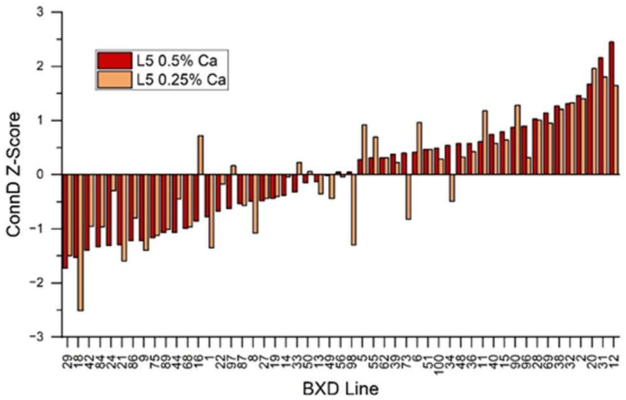


**Low Ca Diet vs RCR**

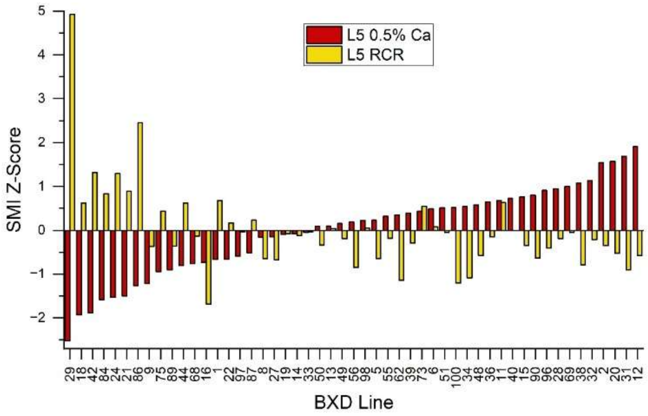


**Low Ca Diet vs RCR**

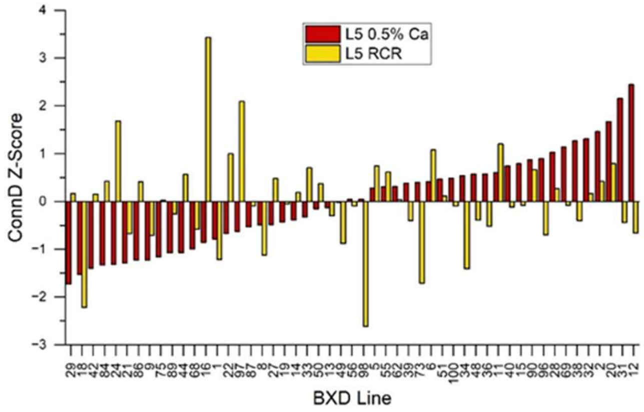


**L5 spine vs Femur**

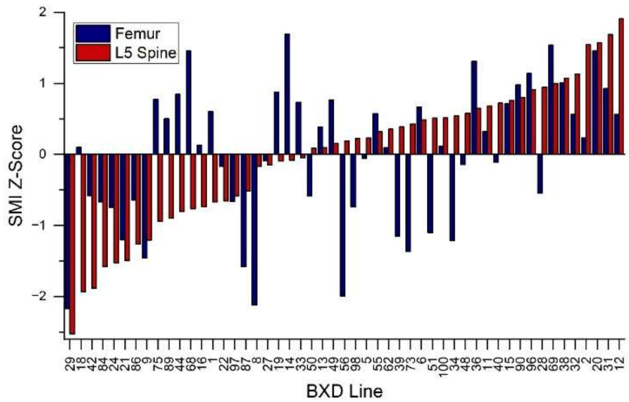


**L5 spine vs Femur**

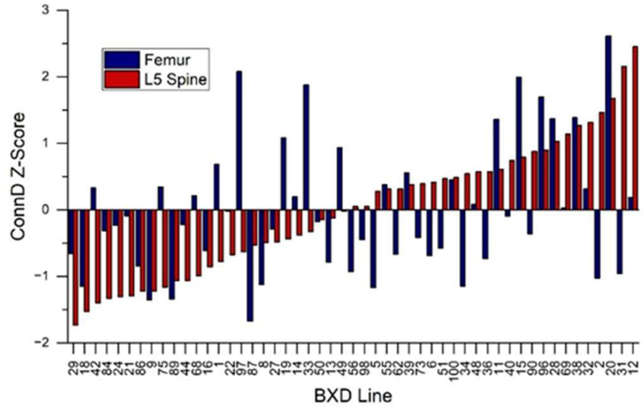

### (G) L5 TMD

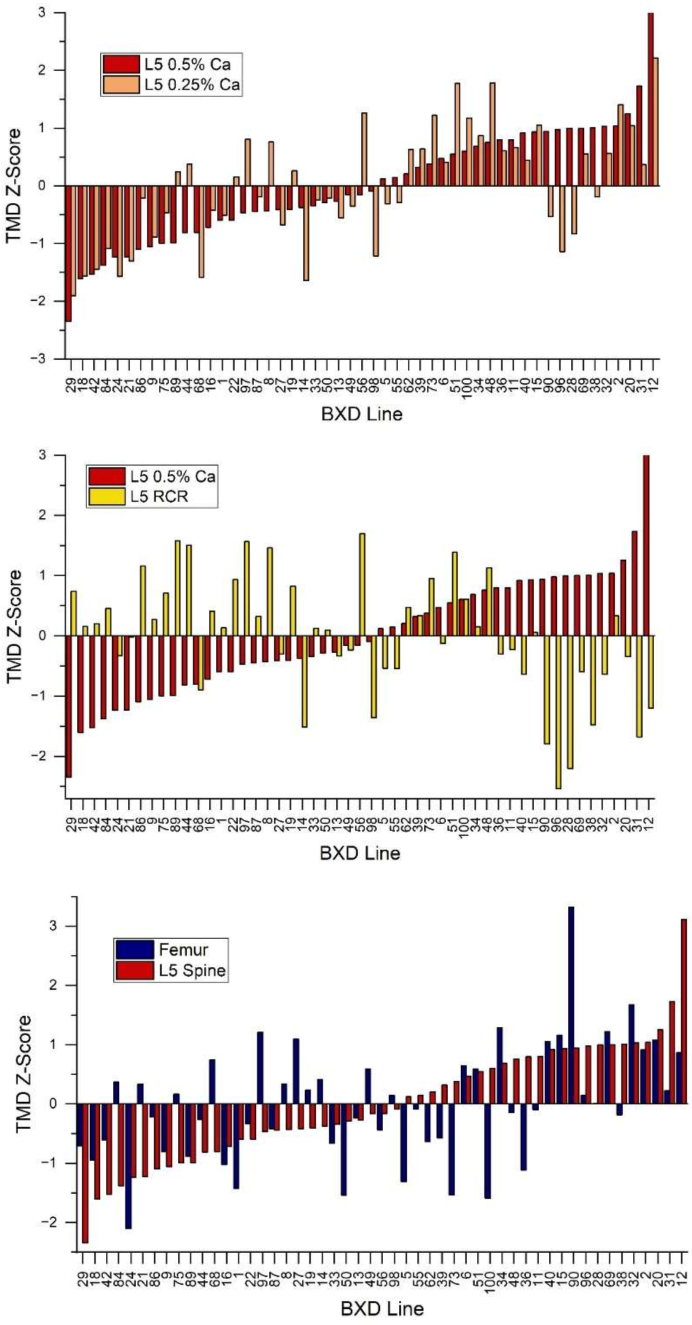
L5 0.5% Ca
L5 0.25% Ca
TMD Z-Score
BXD Line
L5 0.5% Ca
L5 RCR
TMD Z-Score
BXD Line
Femur
L5 Spine
TMD Z-Score
BXD Line

**Supplemental Figure S4.** QTL plots for L5 Spine trabecular bone microCT phenotypes. Values in parentheses are the transformations and the covariate corrections used for the phenotype.

### L5 Spine BVTV

**0.5% Ca** (SQRT(y), BW, FL)

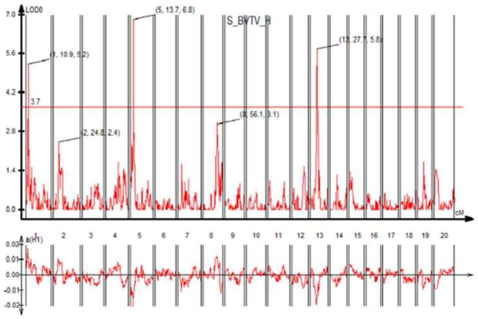

**0.25% Ca** (SQRT(y), BW, FL)

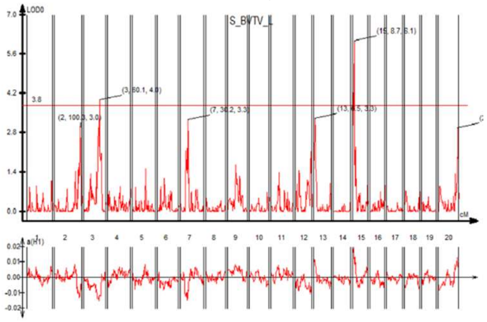

**RCR** (SQRT(y+65), BW, FL)

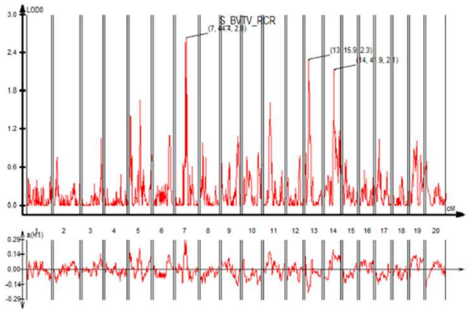

### L5 TbTh

**0.5% Ca** (Log(y), BW, FL)

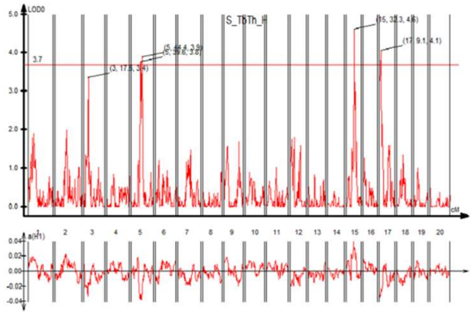

**0.25% Ca** (Log(y), BW, FL)

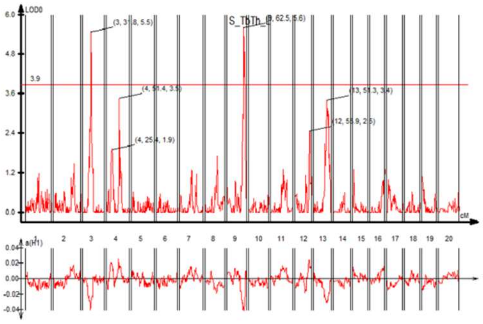

**RCR** (SQRT(y+35), BW, FL)

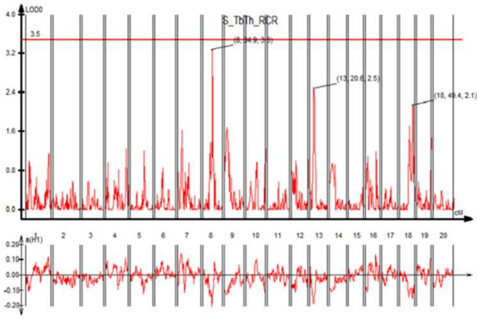

### L5 TbN

**0.5% Ca** (None, BW, FL)

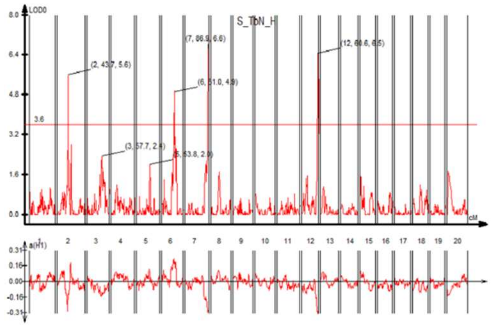

**0.25% Ca** (None, BW, FL)

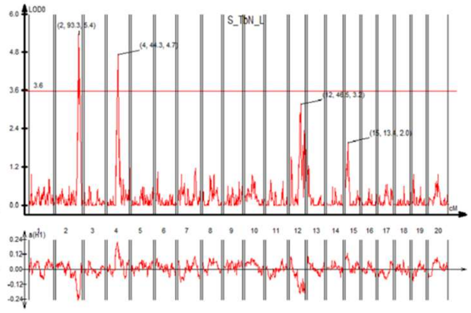

**RCR** (None, BW, FL)

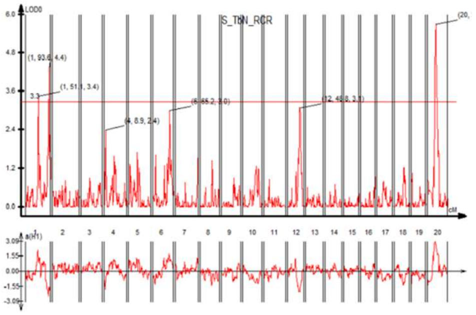

### L5 TbSp

**0.5% Ca** (None, BW, FL)

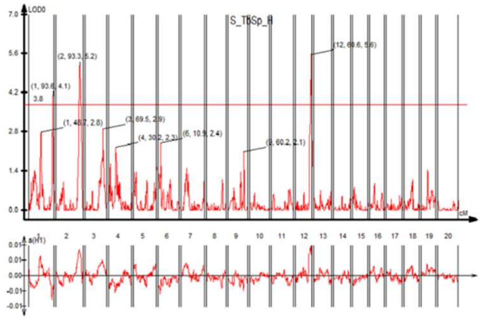

**0.25% Ca** (None, BW, FL)

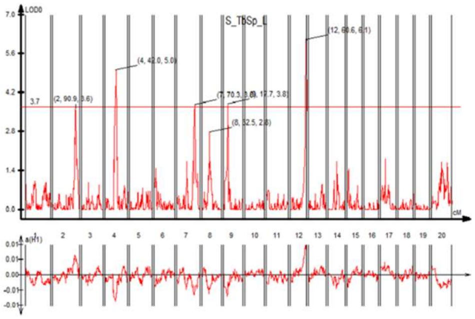

**RCR** (None, BW)

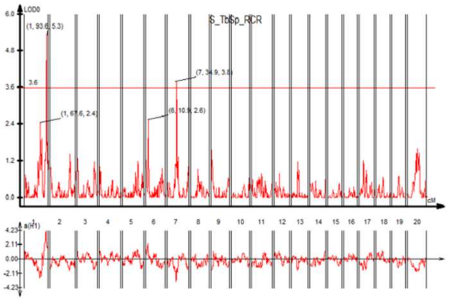

### L5 SMI

**0.5% Ca** (None, BW, FL)

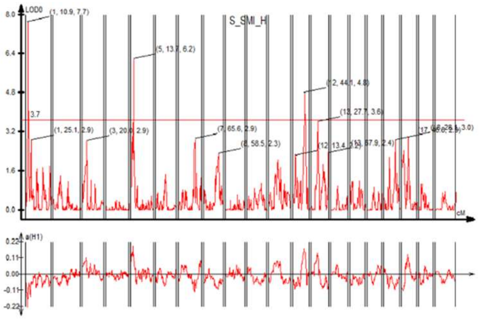

**0.25% Ca** (None, BW, FL)

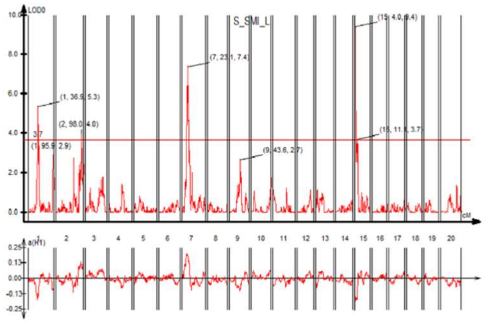

**RCR** (None, FL)

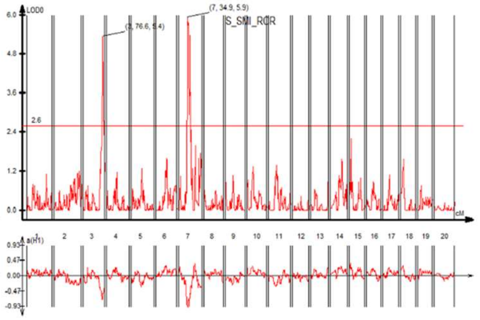

### L5 ConnD

**0.5% Ca** (SQRT(y), BW)

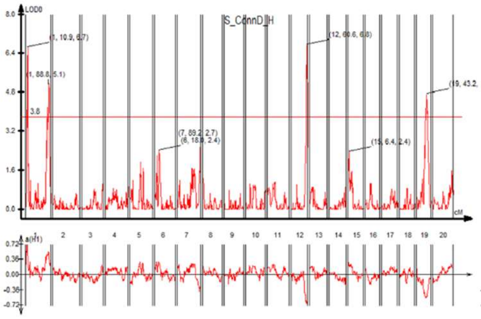

**0.25% Ca** (SQRT(y), BW)

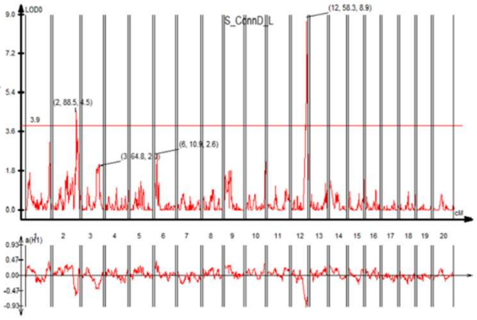

**RCR** (SQRT(y+100), BW)

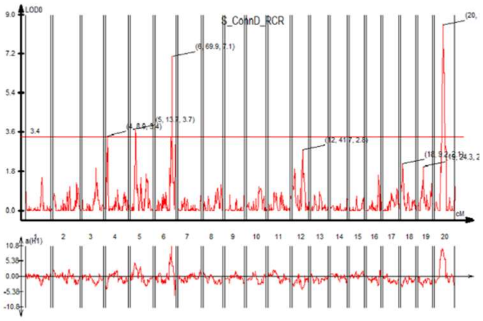

### L5 TMD

**0.5% Ca** (None, None)

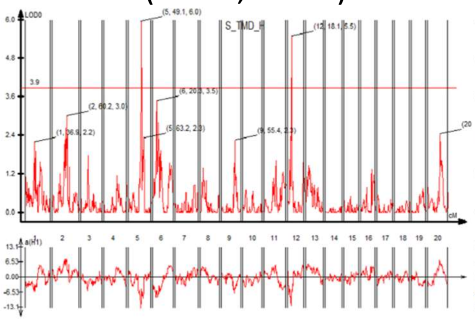

**0.25% Ca** (None, None)

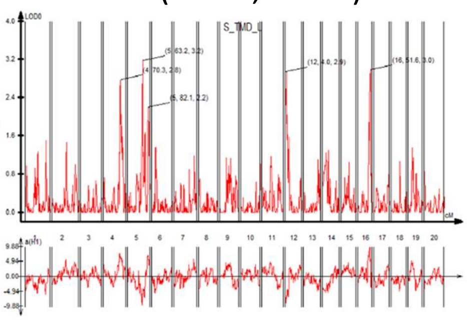

**RCR** (None, None)

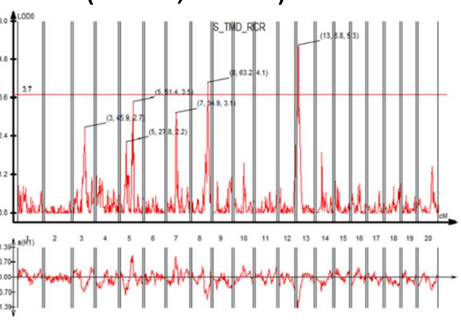
